## Supplemental Figures for "Signaling Diversity Enabled by Rap1-Regulated Plasma Membrane ERK with Distinct Temporal Dynamics"

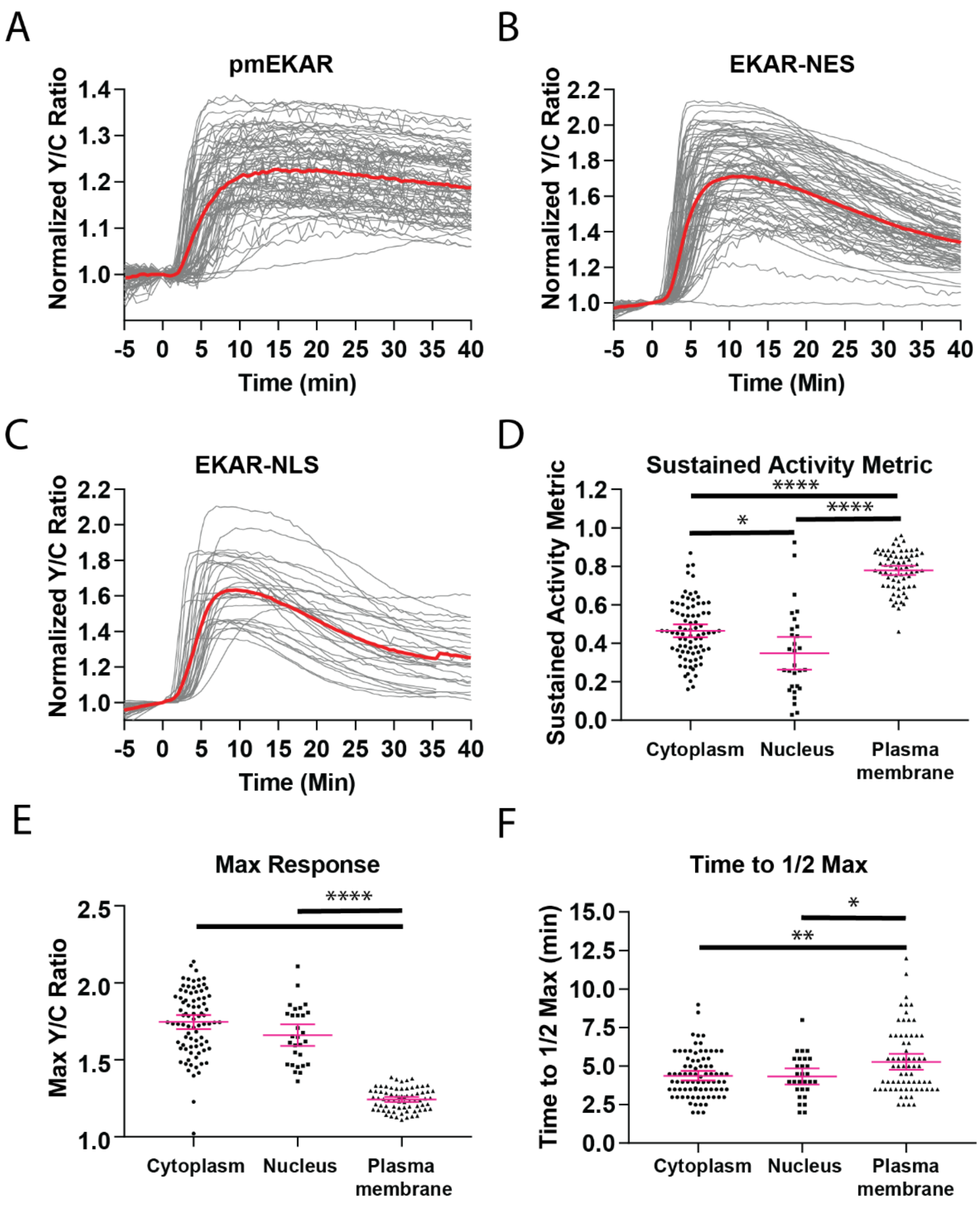


Figure 1 - Supplement 1. A) All cell traces (n = 71) from plasma membrane targeted EKAR4 (pmEKAR4) in cells treated with EGF at time 0. Average shown in red. B) All cell traces (n = 83) of cytosolic EKAR4 (EKAR4-NES) in cell treated with EGF at time 0. C) All cell traces (n = 29) of nuclear-localized EKAR4 (EKAR4-NLS) in cells treated with EGF at time 0. D) Sustained activity metric (SAM40) comparing cytoplasmic, nuclear, and plasma membrane EKAR responses. E) Maximum FRET to donor emission ratio for each EKAR4. F) EKAR4 response time comparisons between cytoplasmic, nuclear, and plasma membrane compartments. The time to ½ maximum responses was recorded and tabulated. Interestingly, cytoplasmic EKAR4 and nuclear EKAR4 responded slightly before pmEKAR4. (Metric comparisons analyzed via 1-way Anova with multiple comparisons, *p<0.05, **p<0.01, ****p<0.0001)

Figure 1-Supplement 2

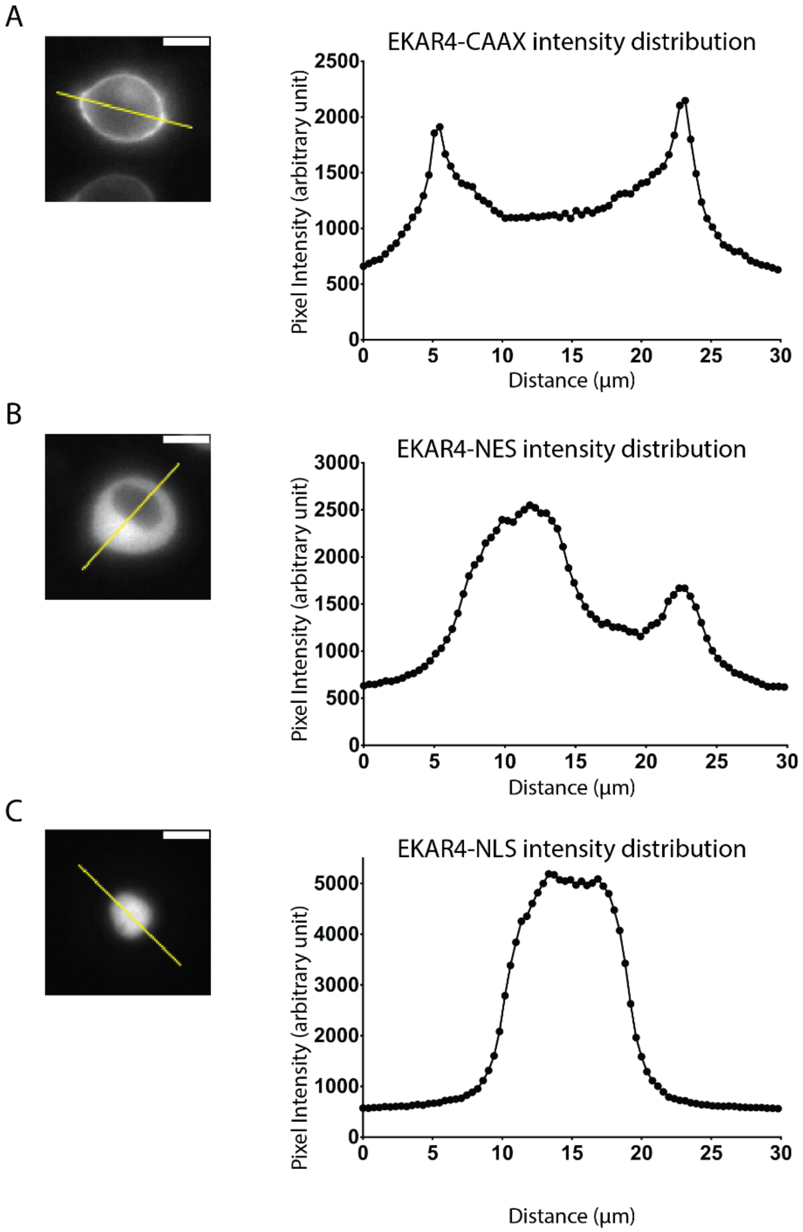


Figure 1 – Supplement 2: Verification of biosensor localization. Representative cells expressing either pmEKAR4 (A), EKAR4-NES (B), or EKAR4-NLS (C) were analyzed by measuring fluorescence intensity of the donor (ECFP) across the indicated line. Scale bar indicates 10 µm.

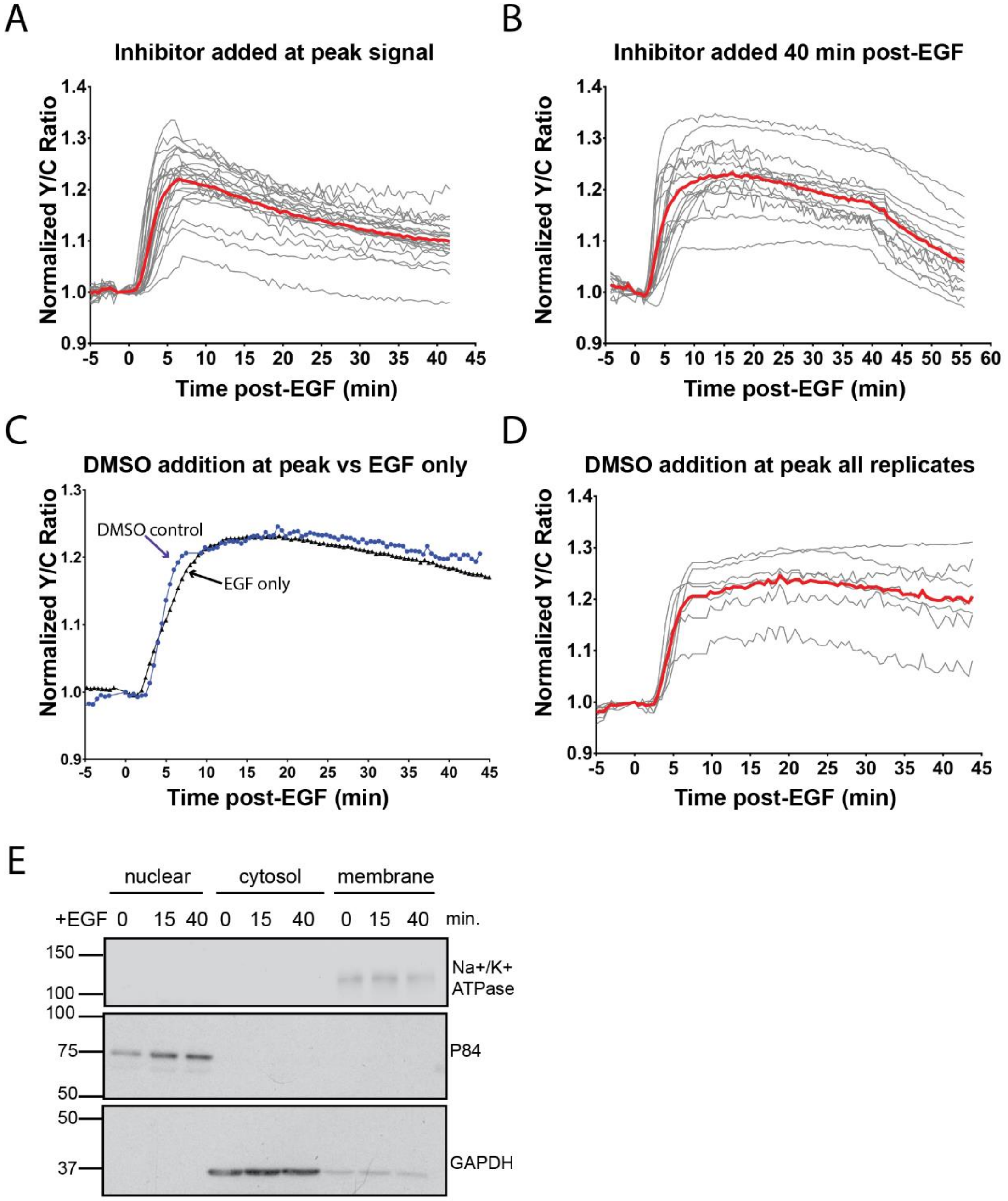


Figure 2 – Supplement 1: A) All Traces (n=22) of pmERK activity in cells treated with 10μM SCH772984 at peak of ERK response to EGF. B) All traces of pmERK activity in cells treated with 10μM SCH772984 40 minutes after EGF addition (n=17). C) Average traces of EGF only (black trace) and DMSO-treated at peak activation (blue trace) of pmEKAR response to EGF. D) All pmEKAR traces of DMSO-treatment at peak EGF response (n=7). E. Verification of successful cellular fractionation. $Na^+/K^+$ ATPase indicates presence of plasma membrane, P84 indicates presence of nucleus, and GAPDH indicates presence of cytosol.

Figure 3 – Figure supplement 1

A

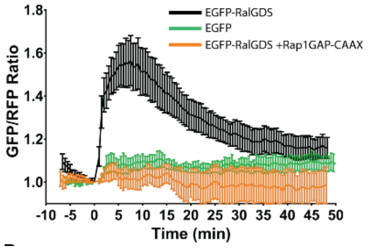


Figure 3 Supplement 1 - Cells expressing either a plasma membrane targeted mCherry-CAAX (mCh) in combination with either EGFP-RalGDS (black curve, n=11 cells), EGFP only (green curve, n=3 cells), or EGFP-RalGDS with Rap1GAP overexpression (orange curve, n=5 cells) were analyzed using TIRF microscopy to track EGFP translocation to the basal membrane, indicating the formation of GTP-bound Rap1 at the plasma membrane.

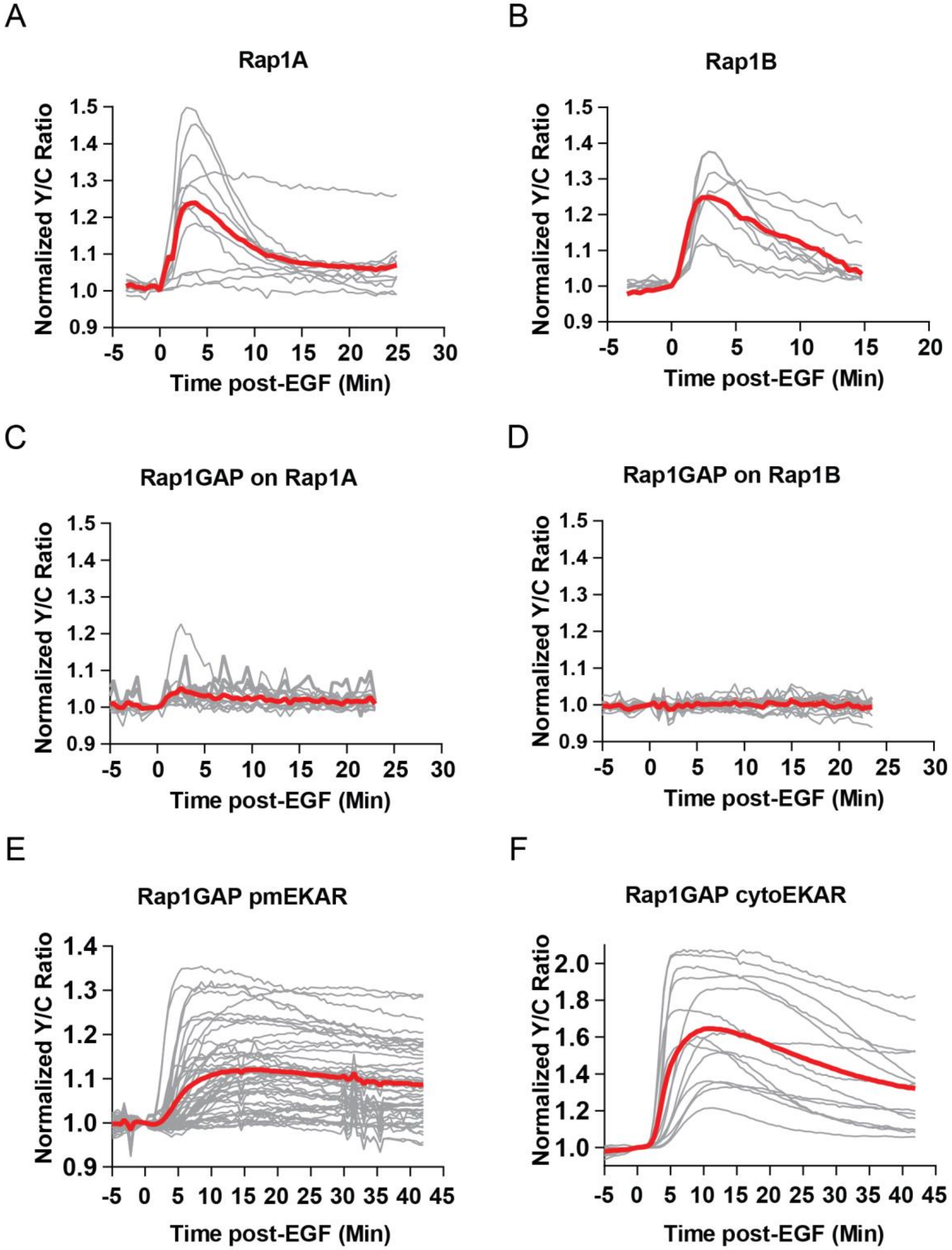


Figure 3 – Supplement 2. A) Rap1A FLARE responses of all replicates, average shown in red (n=11). B) Rap1B FLARE responses of all replicates with average shown in red (n=9). C) Effect of Rap1GAP expression on Rap1A FLARE response, all replicates (n=14). D) Effect of Rap1GAP on Rap1B Flare response, all replicates (n=15). E) Effect of Rap1GAP on pmEKAR responses, all replicates (n=40). F) Effect of Rap1GAP on cytoEKAR responses, all replicates (n=15).

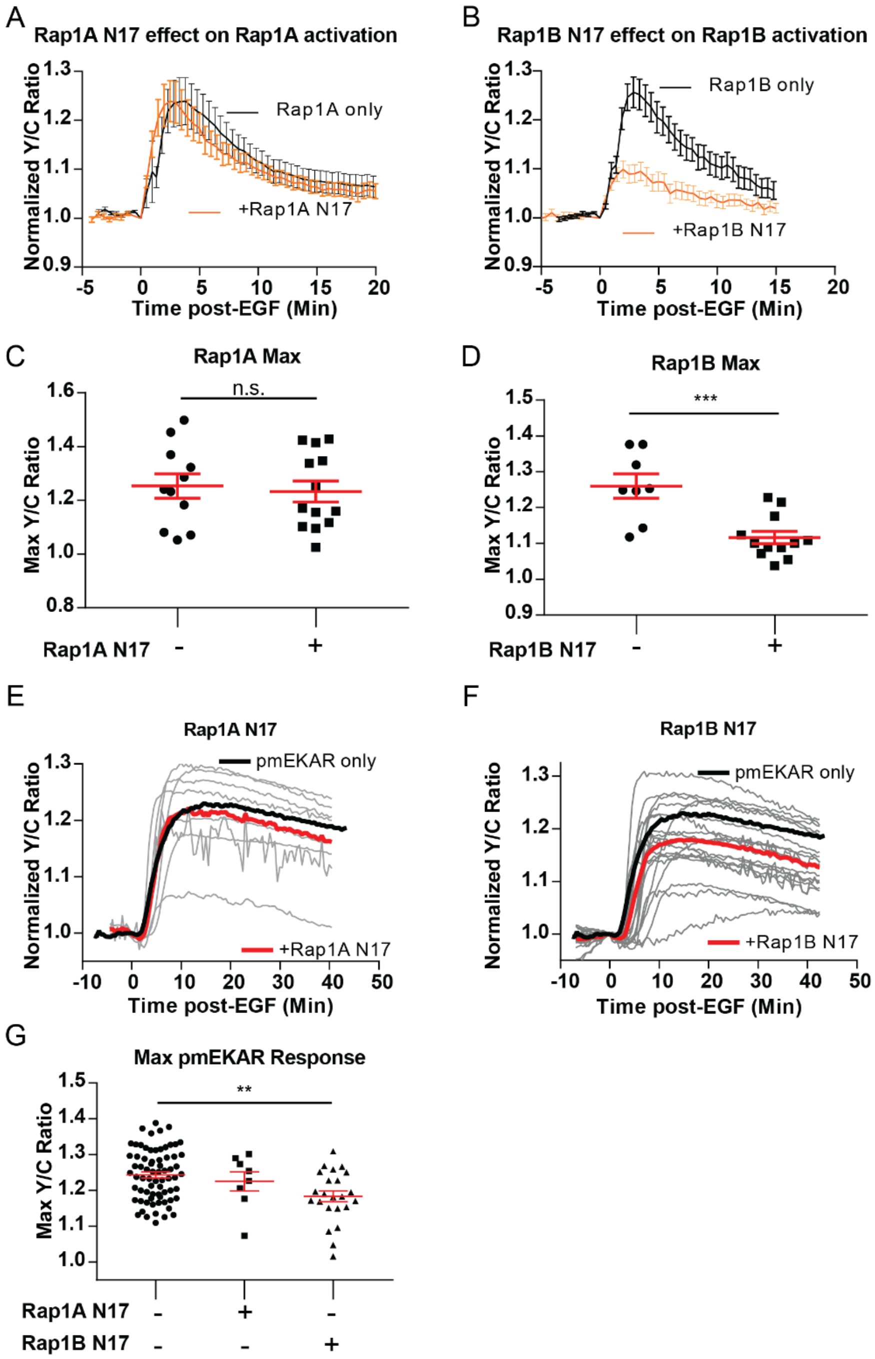


Figure 3 Supplement 3 – Dominant Negative Rap1 isoforms on EGF-induced Rap1 and pmERK activation. Rap1A N17 expression had no effect on Rap1A response (A, C) or pmERK response to EGF (E, G). Rap1B N17 expression significantly dampened Rap1B response (B, D) and dampened pmERK response (F, G) (n=23). Note, Rap1 DN expression resulted in similar trends as Rap1GAP expression, but the effect had more cell-to-cell variability. (***p<0.001, student's t-test; **p<0.01, 1-Way Anova comparison to no dominant negative control)

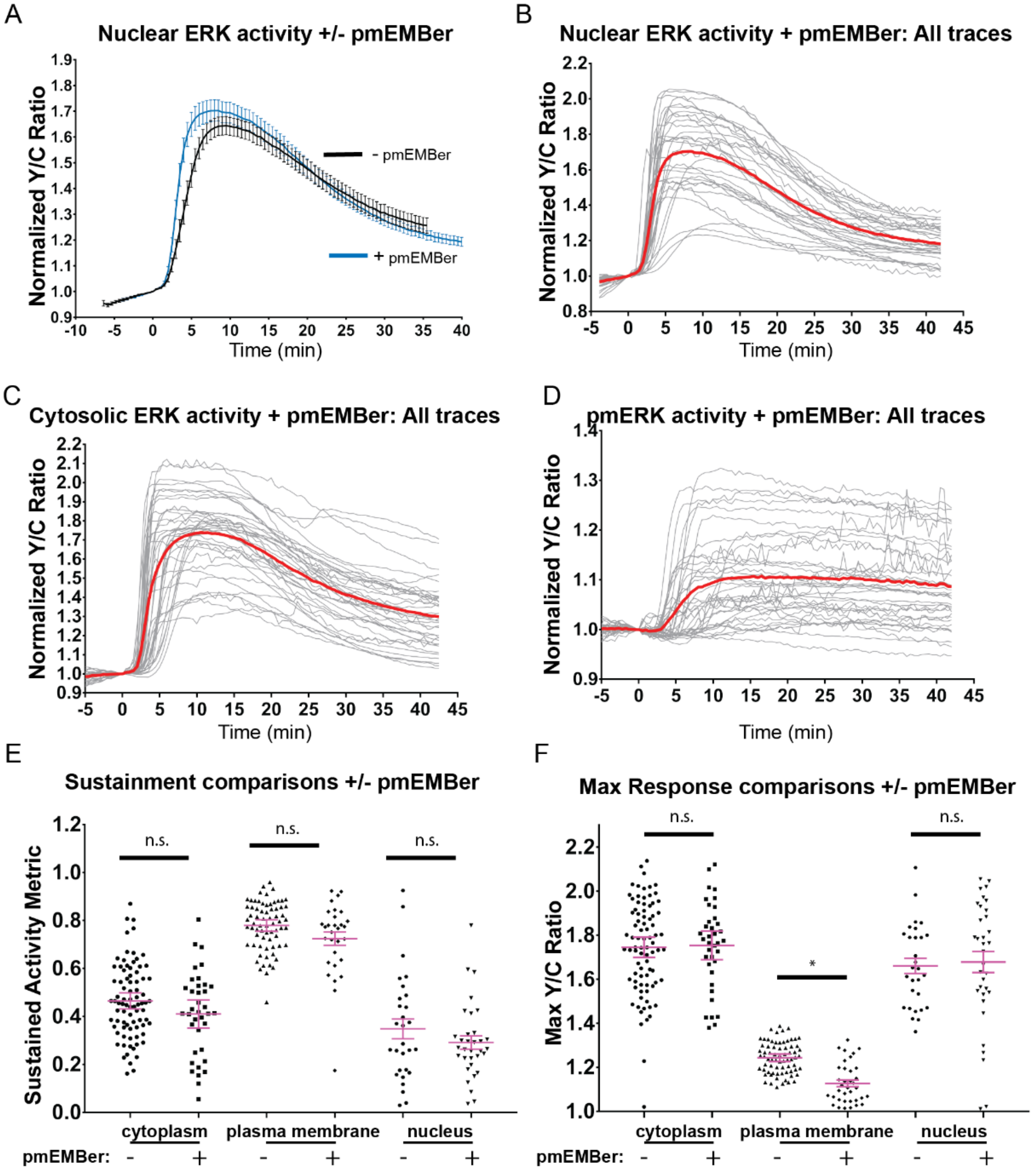


Figure 4 Supplement 1: A) Effect of pmEMBer expression on nuclear ERK response to EGF. B) All traces of nuclear ERK activation in response to EGF in cells expressing pmEMBer (n=32). C) All traces of cytoplasmic ERK activation in response to EGF in cells expressing pmEMBer (n=75). D) All traces of plasma membrane localized ERK activation in response to EGF in cells expressing pmEMBer (n=36). E) Effect of pmEMBer expression on sustainment of ERK responses, measured by SAM40 metric (*Equation 1*). Each condition was compared to every other condition using 1-way ANOVA with multiple comparisons. Each location (plasma membrane, cytoplasm nucleus) showed no significant change in persistence of ERK activity. F) Effect of pmEMBer on maximum response (Y/C ratio) at the cytoplasm, plasma membrane, and nucleus. Each condition was compared to every other condition using 1-way ANOVA with multiple comparisons. (n.s. = not significant, *p<0.05.)

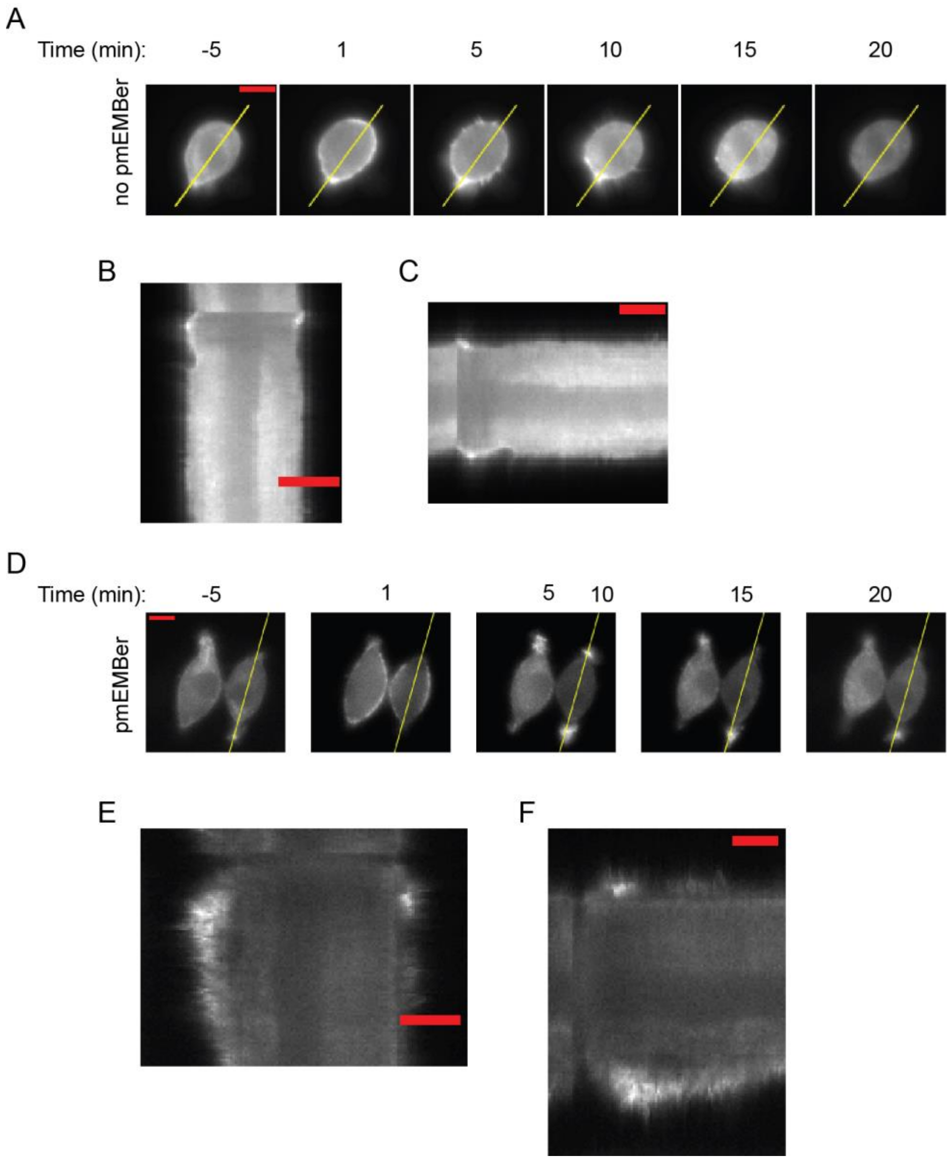


Figure 4 Supplement 2 – Examples of cell protrusion measurements. A, D) Kymograph of indicated area in was obtained along the major axis to quantitate cell protrusions. B, E) Kymographs of cell shown in (A) and (D), respectively. Scale bar represents 10 µm. C, F) Kymographs from (B or E) were rotated so that the horizontal axis represents time. Scale bar represents 600 seconds (10 minutes.)

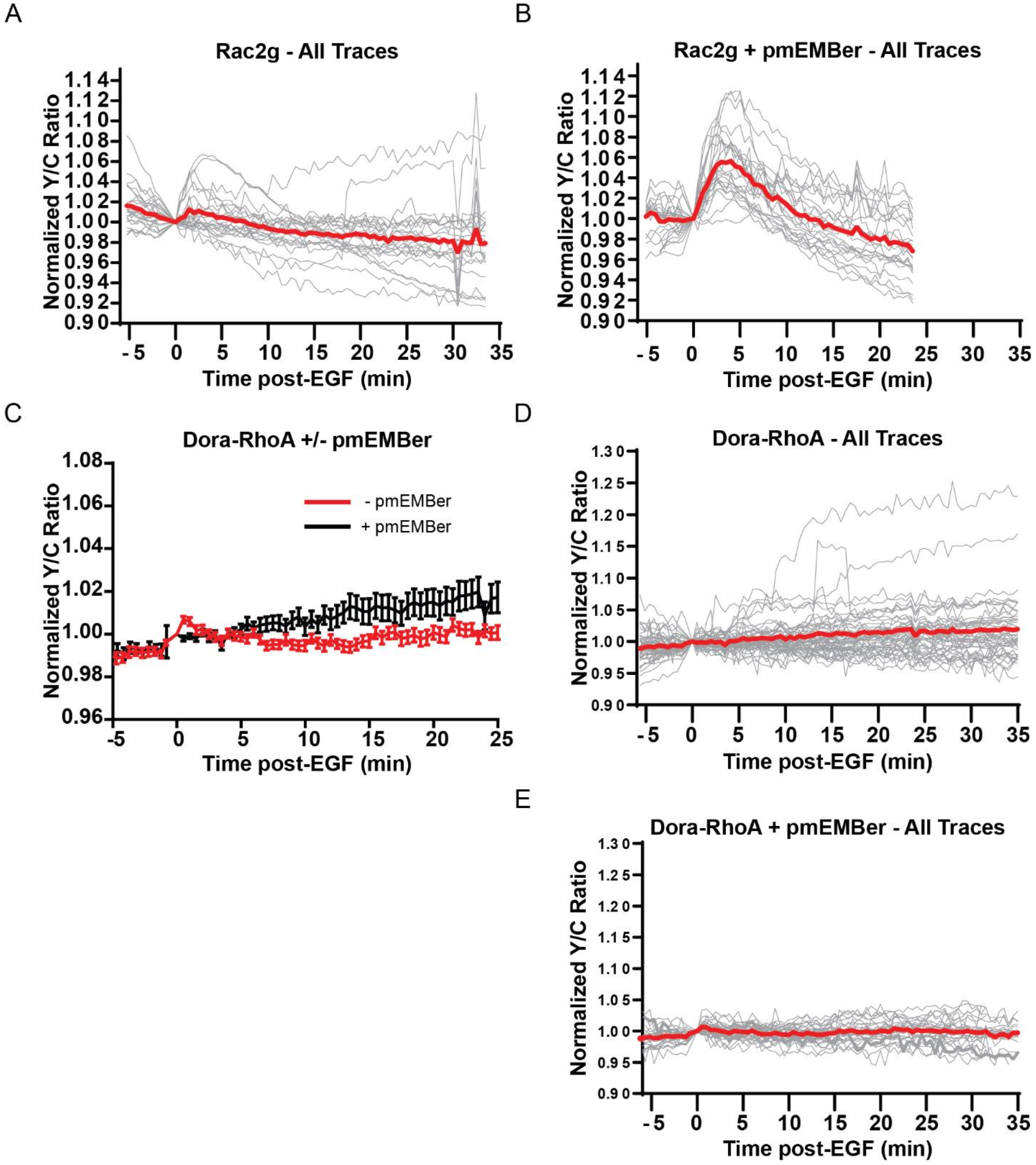


Figure 4 Supplement 3 – A) All traces (n=19) of the effect of EGF on Rac1 measured by the Rac-2G biosensor. Average displayed as red trace. B) All Traces of Rac1 activity in presence of pmEMBer, n=24. C) Average traces comparing RhoA activation in response to EGF with or without pmEMBer expression. D) All traces of RhoA activation in cells expressing mCh-CAAX control (n=50). E) All traces of RhoA activation in cells expressing pmEMBer (n=30).
